## Supplementary Figure 1 for "Predictive learning rules generate a cortical-like replay of probabilistic sensory experiences"

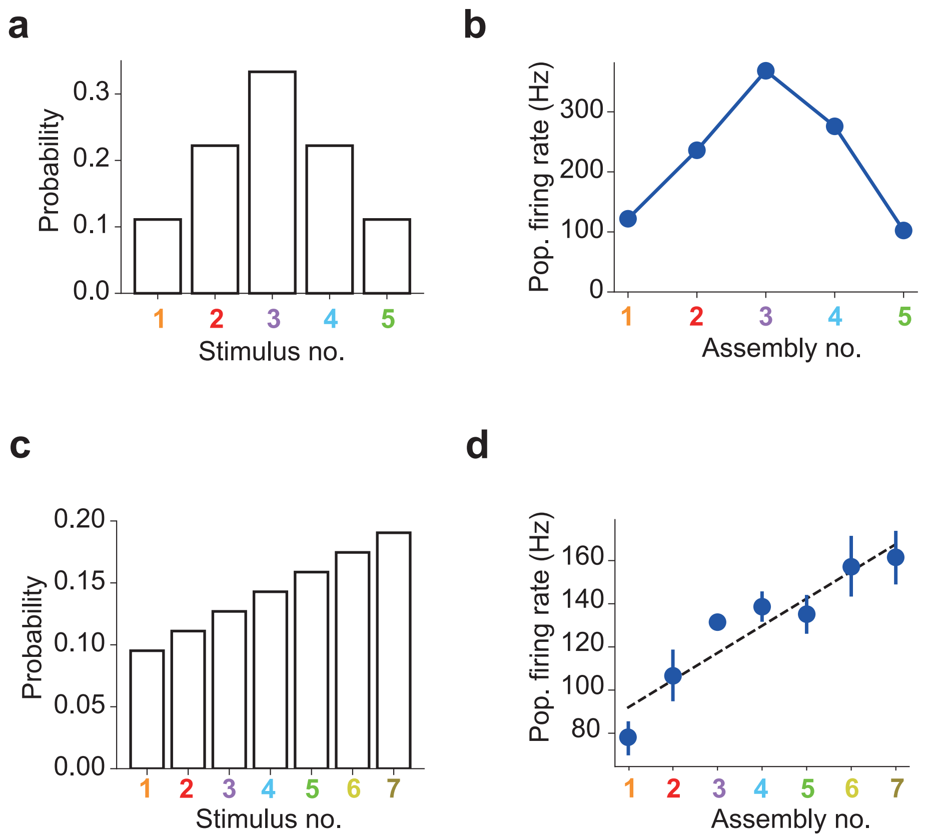


**Supplementary Figure 1. Prior encoding by the nDL model.** As in Fig. 3f and 3g, the nDL model was trained with a set of stimuli. (a) The five stimuli occurred with different probabilities during training. (b) The spontaneously replayed cell-assemblies exhibited population firing rates proportional to the occurrence probabilities of the corresponding stimuli. (c) Similar as in a, but with seven stimuli. (d) The spontaneous population activites of seven assemblies are shown. The activities were propotional to the occurrence probabilities of stimuli shown in c. Error bars show SDs over five independent simulations. A dashed line is a regression line.
