## Supplementary Figure 2 for "Predictive learning rules generate a cortical-like replay of probabilistic sensory experiences"

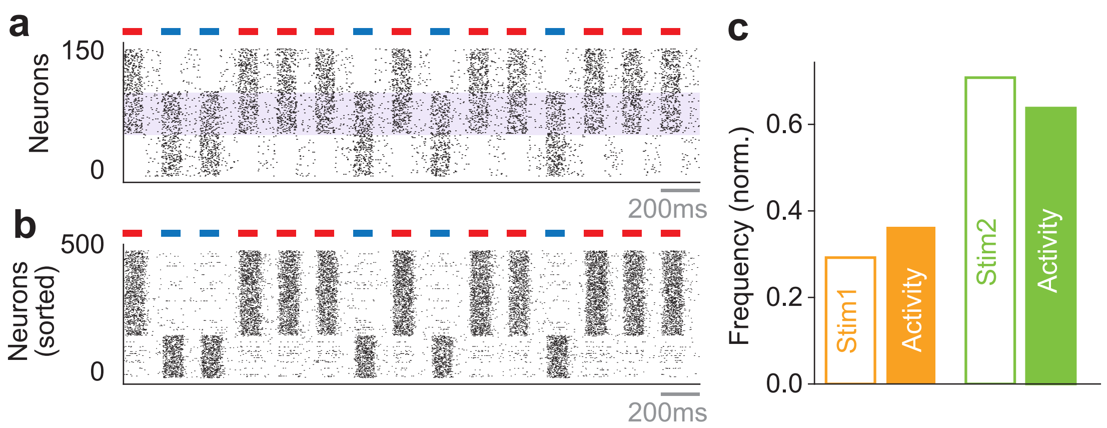


**Supplementary Figure 2. Learning occurence probabilities of overlapped input patterns.** (a) Two input patterns were presented with 30% (blue) and 70% (red) probabilities of which the 50% of input neurons were shared (purple horizontal area). (c) The ratio of the activities of two learned assemblies in a spontaneous activity showed a strong similarity to the stimulus probabilities.
