## Supplementary Figure 3 for "Predictive learning rules generate a cortical-like replay of probabilistic sensory experiences"

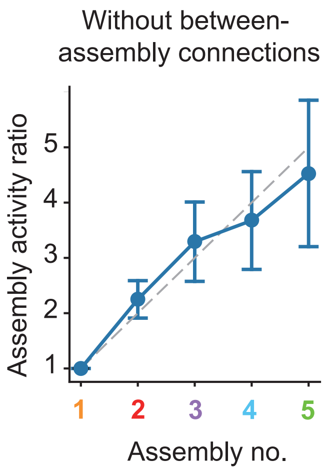


**Supplementary Figure 3. WIthin-assembly connections encode the peobability structures.** The mean ratios of spontaneous population firing rates without between-assembly connections are shown. The connections were removed after the network leaned to encode the stimulus probabilities shown in Fig. 3f. Error bars indicate the SDs over five trials.
