## Supplementary Figure 4 for "Predictive learning rules generate a cortical-like replay of probabilistic sensory experiences"

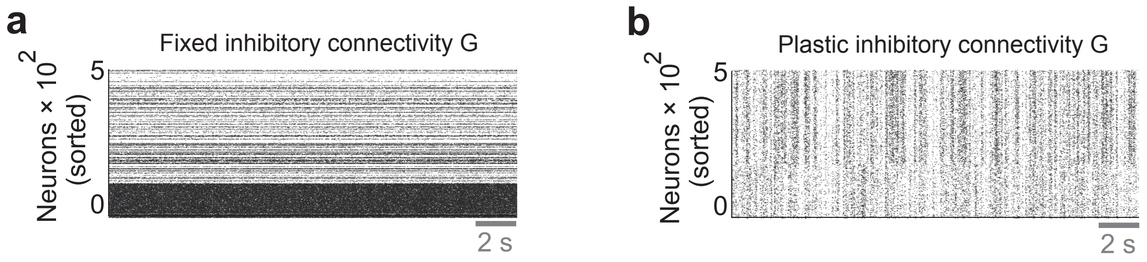


**Supplementary Figure 4. Inhibitory plasticity during learning is necessary to stabilize spontaneous activity.** (a) Spontaneous activity of learned network with non-plastic inhibitory connections during learning. (b) Same as in a, but with plastic inhibitory connections.
