## Supplementary Figure 5 for "Predictive learning rules generate a cortical-like replay of probabilistic sensory experiences"

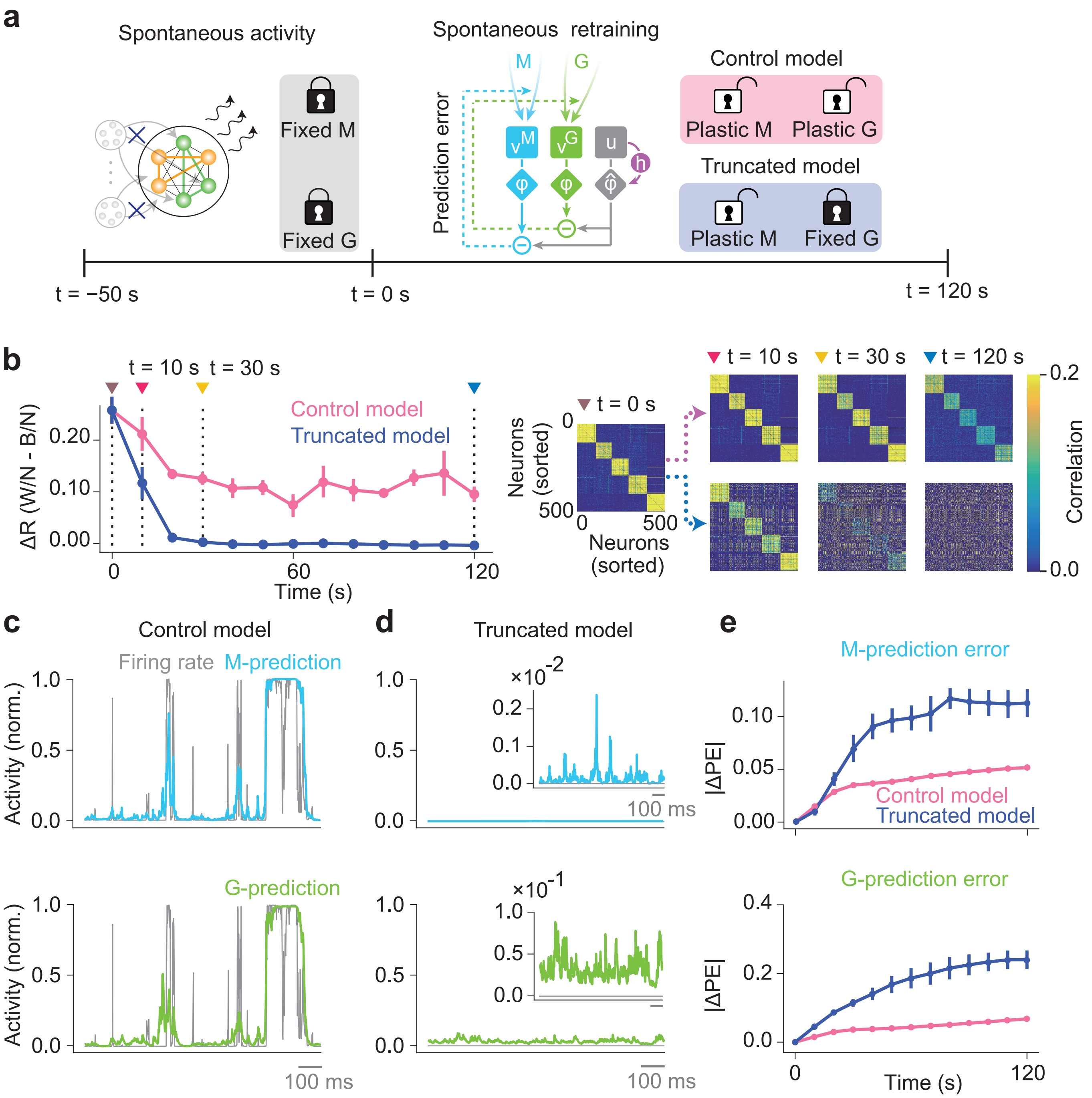


**Supplementary Figure 5. Crucial roles of inhibitory plasticity in prior learning.** We first trained the network models with five external stimuli. (a) Then, we terminated the stimuli at -50 sec and waited until 0 sec for the recovery of network activity through the renormalization process (Eq. 10) with all plasticity rules turned off. We turned on the plasticity of M  at time 0 sec. We kept the plasticity of G turned off in the truncated model (blue), while we turned on the G-plasticity in the control model (magenta). (b) *left*, The time evolution of the difference between the average within-assembly coherence  and the average between-assembly coherence was plotted for the control (magenta) and truncated (blue) models. Larger differences imply more robust cell assemblies. Error bars indicate the SDs over five trials. *right*, Activity coherences between neurons are shown at the indicated times. (c) The time-varying normalized firing rate of a neuron (grey) and the values predicted by recurrent synaptic inputs (top) and lateral inhibition (bottom) are shown for the control model. (d) Similar plots are shown for the truncated model. (e) Changes in prediction errors in the control (magenta) and truncated (blue) models are shown for recurrent synaptic inputs (top) and lateral inhibition (bottom).
