## Supplementary Figure 6 for "Predictive learning rules generate a cortical-like replay of probabilistic sensory experiences"

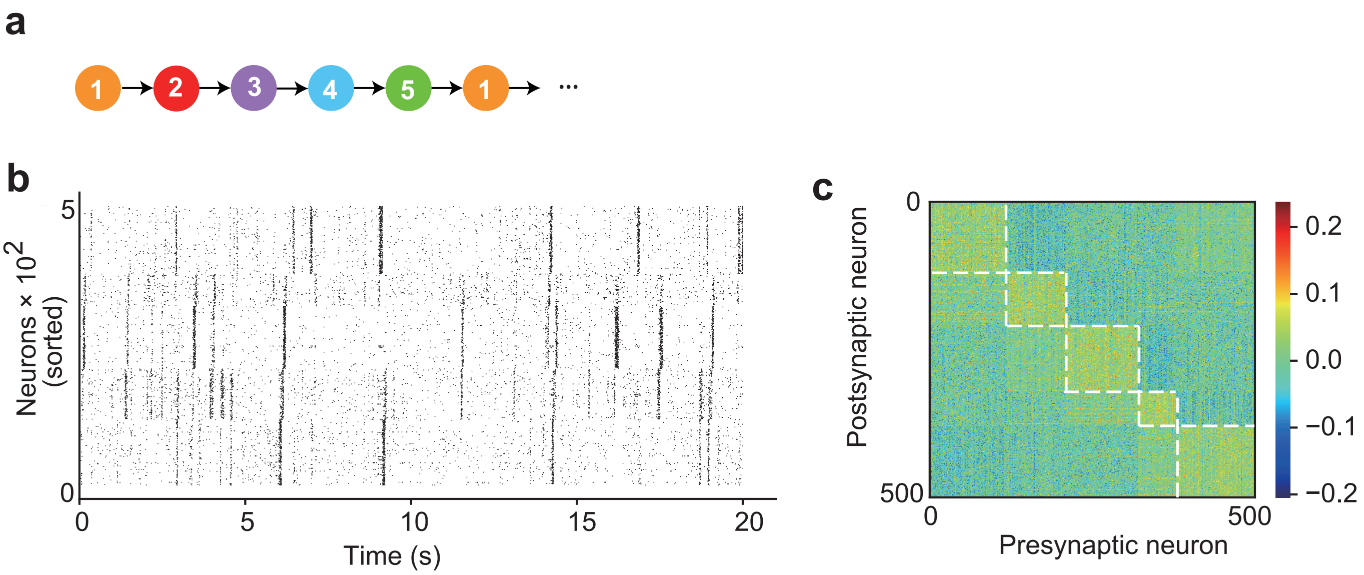


**Supplementary Figure 6. Distinct assembly replay after sequence** (a) The network was trained repetitively with a fixed sequence (i.e., 1-2-3-4-5). (b) An exmaple spontaneous activity after learning. The assemblies are reactivated almost independently. (c) The learned recurrent connection matrix shows stronger intra-assembly and weaker inter-assembly connections.
