## Supplementary Figure 7 for "Predictive learning rules generate a cortical-like replay of probabilistic sensory experiences"

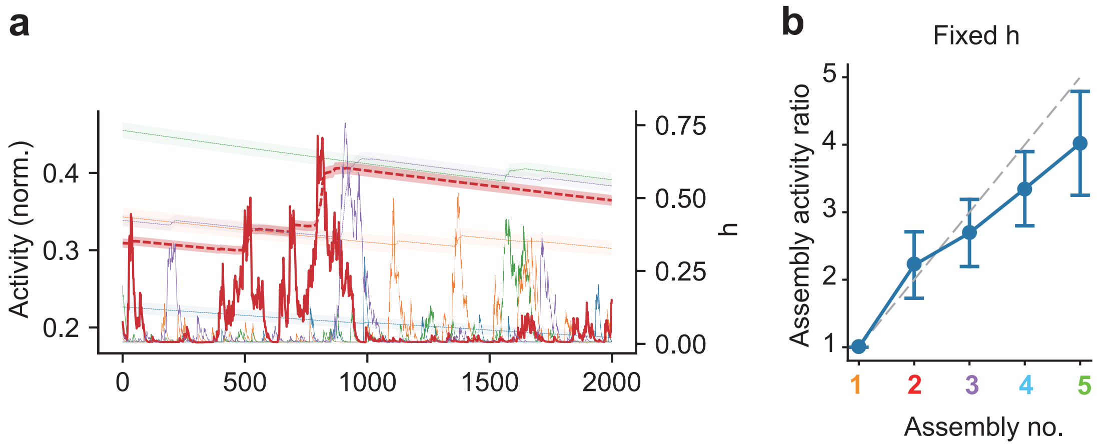


**Supplementary Figure 7. Role of dynamical variable h in spontaneous replay of assemblies.** (a) An example of assembly dynamics (solid) and dynamical variables h (dashed) are shown. Each color refers to one of the five assemblies, and red curves are highlighted for visualization purposes. The dynamical variables show an abrupt increase when the corresponding assembly has peak activity, and a slow decrease otherwise. Note that the dynamics of h corresponding to as-semblies that do not show large activity peaks decay slowly almost everywhere without showing a significant increase (e.g., the green dashed line). Curves show the averaged values over the individual assemblies and shaded areas show the standard error. (b) The mean ratios of spontaneous population firing rates calculated with fixed h variables are shown. The network was first trained to encode the stimulus probabilities shown in Fig. 3f and the h values were then fixed during spontaneous activity. The ratios capture the increasing tendency of the true probability distribution in Fig. 3f with degraded accuracies, especially in assemblies 4 and 5. Error bars indicate the SDs over five trials.
