## Supplementary Figure 8 for "Predictive learning rules generate a cortical-like replay of probabilistic sensory experiences"

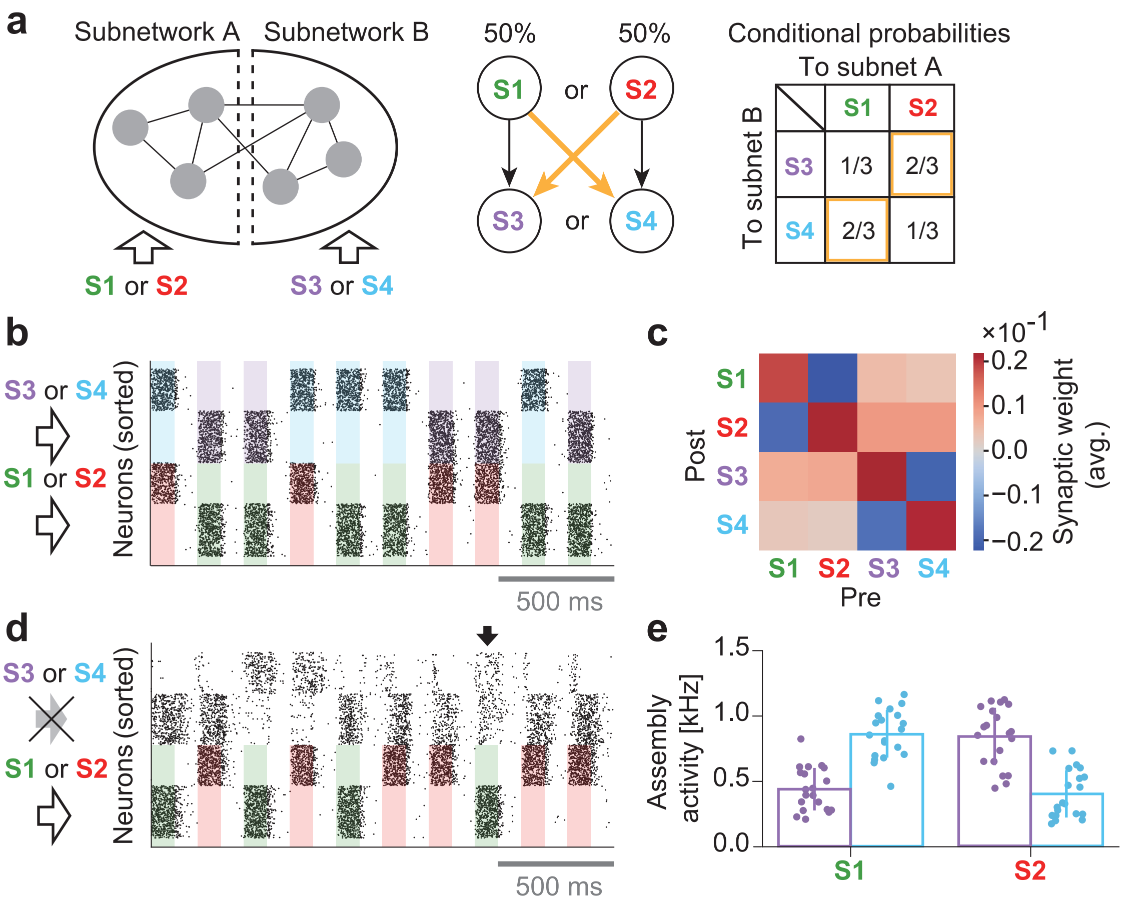


**Supplementary Figure 8. Learning of multivariate priors with assemblies.** (a) Network neurons were separated into two populations receiving different groups of feedforward inputs (left). Subnetwork A received stimuli 1 (S1) and 2 (S2), each presented one at a time with probability 1/2. Subnetwork B received stimuli 3 (S3) or 4 (S4) exclusively when subnetwork A was also stimulated. S3 or S4 was sampled at each presentation according to the probability distribution conditioned on the stimulus presented to subnetwork A (middle and right). (b) Raster plot of evoked activity after training. Each subnetwork formed two assemblies responding to different preferred stimuli. Shaded areas with four colors indicate the duration of stimuli given to the two subnetworks. (c) The connection matrix self-organized among the cell assemblies is shown. (d) The activities of the four assemblies in the presence of S1 and S2 but not S3 and S4 are shown. Despite the absence of stimuli, subnetwork B replayed the assemblies encoding S3 and S4 when subnetwork A was activated by S1 or S2. (e) Activities of assemblies 3 and 4 in subnetwork B varied with the stimulus presented to subnetwork A. Each data point corresponds to one of 20 independent stimulus presentations. Error bars represent SDs.
