## Supplementary Figure 9 for "Predictive learning rules generate a cortical-like replay of probabilistic sensory experiences"

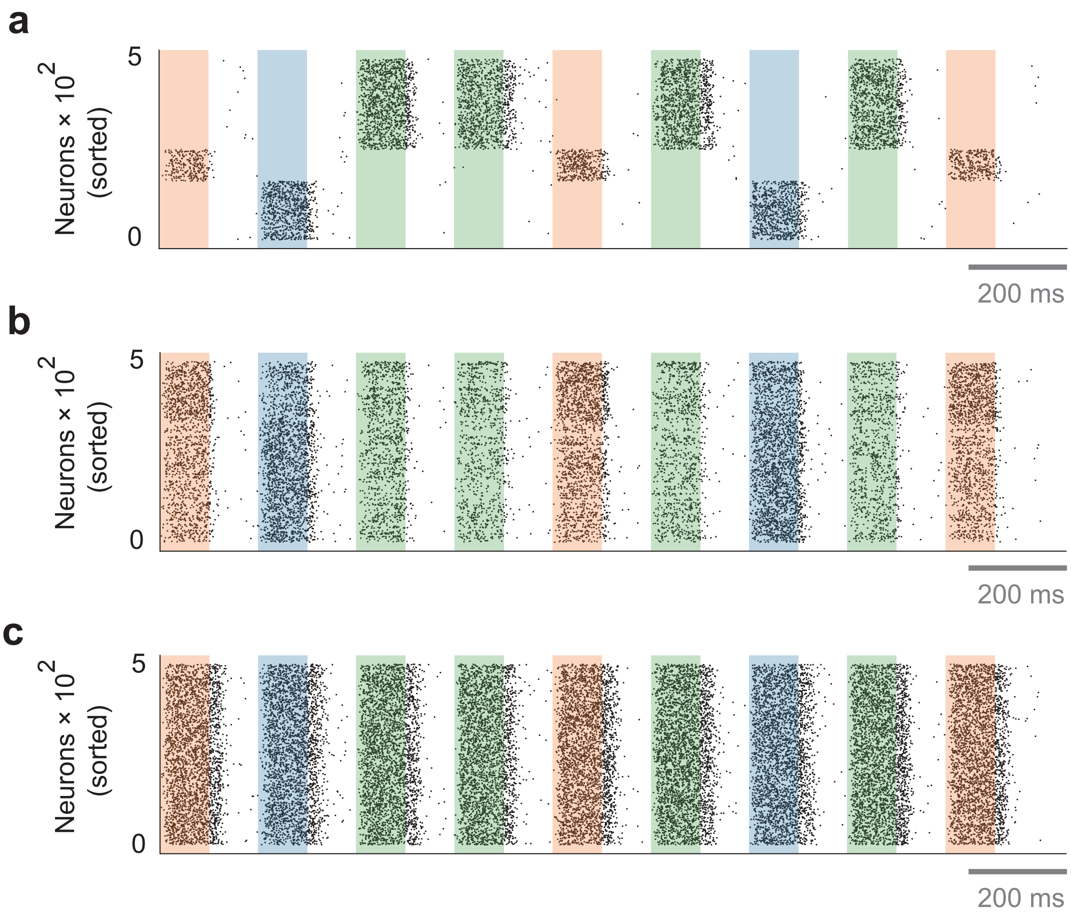


**Supplementary Figure 9. The coexistence of the two inhibitory paths crucial for learning.** (a) A typical spike raster of stimulus-evoked responses is presented for the elaborate network model shown in Fig. 7. (b) A spike raster of stimulus-evoked responses is shown for simulations of the elaborate model without inhibitory path 2. Inhibitory connections were modifiable in path 1. (c) A similar spike raster is presented for simulations of the elaborate model without inhibitory path 1. Inhibitory connections were modifiable in path 2. The results shown in b and c demonstrate that the network model fails to self-organize the cell assemblies encoding the different stimuli when it lacks one of the two inhibitory paths.
